# High-depth resequencing reveals hybrid population and insecticide resistance characteristics of fall armyworm (*Spodoptera frugiperda*) invading China

**DOI:** 10.1101/813154

**Authors:** Lei Zhang, Bo Liu, Weigang Zheng, Conghui Liu, Dandan Zhang, Shengyuan Zhao, Pengjun Xu, Kenneth Wilson, Amy Withers, Christopher M. Jones, Judith A. Smith, Gilson Chipabika, Donald L. Kachigamba, Kiwoong Nam, Emmanuelle d’Alençon, Bei Liu, Xinyue Liang, Minghui Jin, Chao Wu, Swapan Chakrabarty, Xianming Yang, Yuying Jiang, Jie Liu, Xiaolin Liu, Weipeng Quan, Guirong Wang, Wei Fan, Wanqiang Qian, Kongming Wu, Yutao Xiao

## Abstract

The rapid wide-scale spread of fall armyworm (*Spodoptera frugiperda*) has caused serious crop losses globally. However, differences in the genetic background of subpopulations and the mechanisms of rapid adaptation behind the invasion are still not well understood. Here we report a 393.25-M chromosome-level genome assembly of fall armyworm with scaffold N50 of 13.3 M consisting of 23281 annotated protein-coding genes. Genome-wide resequencing of 105 samples from 16 provinces in China revealed that the fall armyworm population comprises a complex inter-strain hybrid, mainly with the corn-strain genetic background and less of the rice-strain genetic background, which highlights the inaccuracy of strain identification using mitochondrial or *Tpi* genes. An analysis of genes related to pesticide- and Bt-resistance showed that the risk of fall armyworm developing resistance to conventional pesticides is very high, while remaining currently susceptible to Bt toxins. Laboratory bioassay results showed that insects invading China carry resistance to organophosphate and pyrethroid pesticides, but are sensitive to genetically modified maize expressing Cry1Ab in field experiments. Additionally, we found that two mitochondrial fragments are inserted into the nuclear genome, and the insertion event occurred after the differentiation of the two strains. This study represents a valuable advancement toward the analysis of genetic differences among subpopulations and improving management strategies for fall armyworm.

## Introduction

The fall armyworm, *Spodoptera frugiperda* (J.E. Smith), is a polyphagous pest that is native to tropical and subtropical America, with a strong capacity for migration and reproduction^1–4^. The pest was first detected in Africa in 2016^5^ and spread to 44 African countries within two years. It was first detected in Asia in July 2018, and so far it has spread to 13 countries in the region (https://gd.eppo.int/taxon/LAPHFR/distribution). Such rapid spread poses a global threat to food production. The strong environmental adaptability of fall armyworm is not only reflected in its polyphagy for a wide range of host plants^6^, but also in its evolution of resistance to chemical pesticides and genetically modified crops expressing *Bacillus thuringiensis* (Bt) toxins^7–16^. Studies have shown that the genes related to detoxification and metabolic processes in the fall armyworm have exhibited obvious expansion^17–18^. In addition, there are two morphologically identical, but genetically distinct, subpopulations or strains of fall armyworm, the rice-strain (R-strain) and the corn-strain (C-strain), which differ in their host plant selection and sex pheromone composition^19–22^. However, there is no absolute mating barrier between the two strains and productive hybridization has been confirmed in both laboratory and field studies^23–24^.

At present, a number of field-evolved resistant populations of fall armyworm have been detected, including resistance to a variety of chemical pesticides and Bt crops, and the level of resistance is increasing^25–29^. The mechanisms of resistance to pesticides are mainly due to variation in receptor genes, such as amino acid mutations in the ryanodine receptor (RyR) (diamide), acetylcholinesterase (AChE) (organophosphate), voltagegated sodium channel (VGSC) (pyrethroids), and so on^30–32^. In addition, the frame-shift mutation resulting in early termination of the *ABCC2* gene, caused by a 2-bp insertion, is linked to resistance to Cry1Fa^33^. Field-evolved strains resistant to Vip3Aa20 were obtained by screening homozygous resistance loci in F_2_ generations in the laboratory^34^. The resistance mechanisms of other Cry toxins from Bt are still unknown. Clarifying the development of pesticide- and Bt-resistance in fall armyworm would be helpful in providing scientific support for the commercialization of genetically modified crops and Bt biopesticides.

Recent studies have indicated that the molecular identification of the two strains of fall armyworm is dependent on which markers are used^35–37^. The early molecular markers based on mitochondrial *Cytochrome Oxidase Subunit I* (*COI*) and Z-chromosome-linked *Triosephosphate isomerase* (*Tpi*) genes failed to accurately assign strain identification^38–41^. The dominant population of fall armyworm invading Africa and Asia were speculated to be hybrid populations of the female R-strain and male C-strain, based on these two molecular markers^42^. In addition, an Africa-specific haplotype, different from those of native Americas, was also reported in African and Chinese samples based on the *Tpi* gene^18, 40^, which makes strain identification and population genetic structure more complicated. Therefore, a genome-wide analysis of the genetic characteristics of invasive fall armyworm is becoming imperative. Although several versions of the fall armyworm genome have now been published^17–18, 43–44^, the different mechanisms directly related to the two strains are unclear, and the debate about strain identification requires further genomic support and explanation. Here we report a chromosome-level genome sequence of a male moth from an inbred fall armyworm strain, representing a C-strain *COI* and an Africa-specific *Tpi* haplotype which was different from the Western Hemisphere (henceforth American) R-strain and C-strain. We also re-sequenced 105 fall armyworm samples from 16 Provinces in China, as well as four samples collected from two African countries (Zambia and Malawi). The genome-wide genetic backgrounds of the invading fall armyworm samples were compared, and the insecticide-resistance risk was assessed based on analysis of potential resistance-related genes. Comparative genomic analyses of these data help to reveal the resistance-related mechanisms and the population genetic characteristics of fall armyworm, which may facilitate its future management.

## Results

### High-quality genome landscape of fall armyworm

Genomic DNA of fall armyworm was extracted from a male moth of an inbred laboratory-reared strain, and sequencing reads were obtained using both PacBio and Illumina technologies and were assembled after filtering out the low-quality and duplicated reads. A total of 24.72 G PacBio long reads and 162.4 G of high-quality Illumina reads were generated, representing an ~460 × coverage of the fall armyworm genome. Through wtdbg2^45^, the final genome was assembled into 777 contigs with size of 393.25 M and contig N50 length of 5.6 M (longest, 18 M), including a complete mitochondrial sequence. The assembled genome size was close to the estimated size of 395 M based on k-mer depth distribution analysis, which was also similar to that of flow cytometry (396 ± 3M) ^17^. After interaction analysis based on a total of 78 G data obtained through Hi-C sequencing, 497 contigs were concatenated to 31 chromosomes with the scaffold N50 of 13.3 M, accounted for 98.67% of total genome length (Supplementary Fig. 1, Supplementary Table 1). By aligning the Illumina data with the assembled fall armyworm genome, the mapping rate and coverage were 98.76% and 99.68% respectively, which showed the accuracy and high integrity of genome assembly. The genome size reported in this study is intermediate between those of previously published fall armyworm versions, but the genome is nearly 140 M smaller than that recently published by Liu et al. (2019). By genome collinearity analysis, the alignment showed that more than 98% of the current assembled genome was consistent with previously published versions (Supplementary Table 2), indicating that the genome integrity of this study was high, and the previous assembled genome with larger size was mainly caused by high heterozygosity of sequenced samples.

By combining homology-based and *de novo* approaches, we identified ~27.18% of repetitive elements in the assembled fall armyworm genome. Among the known repeat families, LINE constituted the most abundant repeat family, representing 8.66% of the repetitive sequences, while LTR was only 1.38% (Supplementary Table 3). To annotate the fall armyworm genome, we performed deep transcriptome sequencing of larvae, pupae, male and female moths, including three different developmental stages, which generated 98.4 G of RNA sequencing data. By combing homologue-based, *ab initio* and transcriptome-based approaches, we predicted 23281 protein-coding genes (gene models) in the fall armyworm genome, which is greater than the number of predicted genes in other lepidopteran genomes that have so far been published^46–51^. The average CDS length is 1476 bp, with 5.39 exons per gene and an average intron size of 1165 bp, which is larger than those in previously published fall armyworm genomes. More than 85.5% of the predicted coding sequences were supported by transcriptome sequencing data. Further assessment of assembly integrity based on Benchmarking Universal Single-Copy Ortholog (BUSCO) analysis shows that the current genome covered 98.2% complete BUSCO genes (Supplementary Table 4), indicating the high accuracy of the gene predictions.

Comparative analysis of orthogroups of nine Lepidoptera species and *Drosophila melanogaster* (Diptera) were performed. Among them, 17571 groups were found in the current genome of fall armyworm, mainly in the number of other and unassigned genes. Compared with *Spodoptera litura*, *S. frugiperda* has more species-specific genes, and the number of unassigned genes is much greater than that of *S. litura* (Fig. 1a). Phylogenomic analyses of the ten species were conducted using 1571 single-copy genes. As shown in Figure 1a, the phylogenetic tree reflects the taxonomic relationship and phylogenetic status of the different species. Three species of Noctuidae, including *S. frugiperda*, formed one group, which then clustered with *Bombyx mori* (Bombycidae) and *Manduca sexta* (Sphingidae). Two butterflies *Danaus plexippus* and *Heliconius melpomene* (both Nymphalidae) clustered together as an outer branch, while *Plutella xylostella* (Plutellidae) is the outermost branch of Lepidoptera (Fig. 1a).

**Figure 1.**
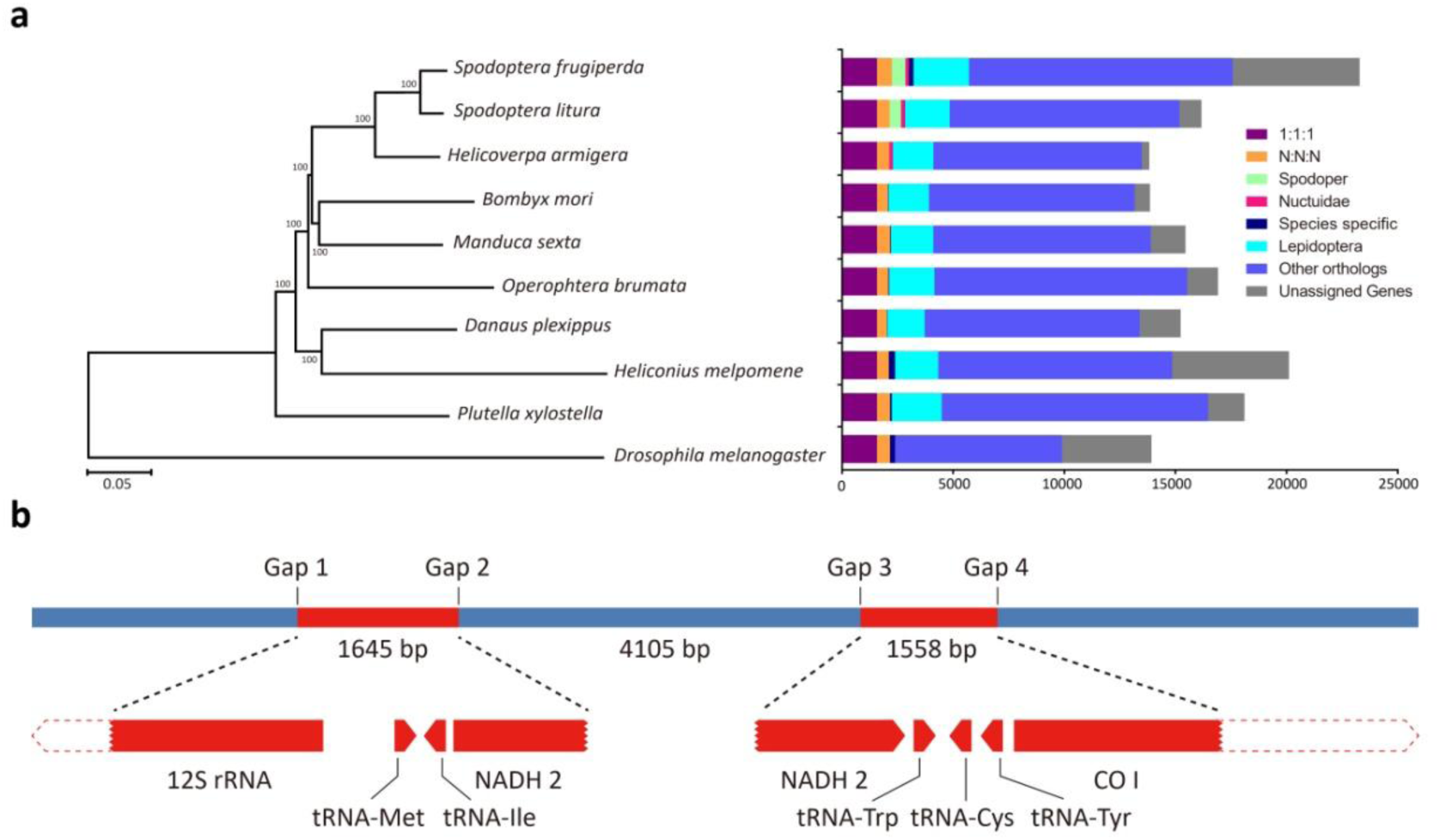
Phylogenetic relationships and schematic map of mitochondrial insertion. a) Phylogenetic tree and genomic comparison of 10 species of Lepidoptera and Diptera. *Drosophila melanogaster* was used as an outgroup and bootstrap value was set as 1000, 1:1:1 include the common orthologs with the same number of copies in different species, N:N:N include the common orthologs with different copy numbers in different species, other orthologs include the unclassified orthologs, and unassigned genes include the genes that cannot be clustered into known gene families. b) A schematic map of two mitochondrial fragments inserted into the nuclear genome, *NADH2* gene was separated by a 4105-bp fragment, and both two inserted mitochondrial fragments were identical with C-strain genotype.

### Insertion of mitochondrial fragments into nuclear genome in a recent evolution event

We found that two mitochondrial fragments, with sequence lengths of 1.5 K (partial *COI* gene and *NADH2* gene) and 1.6 K (partial *NADH*2 gene and 12S rRNA gene), were inserted into the nuclear genome, separated by a 4 K segment of the nuclear genome (Fig. 1b). The total length of a ~7.3K fragment, including two inserted fragments, was supported by more than 28 original reads of PacBio data. The lengths of all 28 original reads were longer than 20 K and completely covered the 7.3 K fragment. However, the two insertions were not found in other published fall armyworm genomes. In order to verify the accuracy of this result, we designed four primers based on flanking sequences of four connection points, and the results of PCR amplification confirmed the existence of the insertion. In addition, four primers were applied to detect the insertion in 173 fall armyworm samples collected from different regions of China and it was found that the insertion was only present in 26.01% of all samples (Supplementary Table 5). At the same time, the resequencing Illumina data of 109 fall armyworm samples in this study also showed that there were varying numbers of reads covering the four junction points in 30 samples, and the percentage of samples with inserted reads was 28.5% (Supplementary Table 6). Both the PCR and resequencing results showed that the insertion was not present in all samples, suggesting that it was a late evolution event.

Moreover, the genotype of the two inserted mitochondrial fragments were identical with those of the C-strain, indicating that the insertion occurred after the differentiation of the R- and C-strains, and was more likely to be a random recombination event between the two strains. Further analysis indicated that two mitochondrial fragments were inserted into the intron region of lysine-specific demethylase 3 B (Kdm3B) gene, which is not likely to affect the expression of the gene. The inserted partial *COI* and *NADH2* gene fragments were also not tend to express and thus play a mitochondrial related function. However, such two fragments could be used to develop markers to identify specific populations and used for further evolutionary events of fall armyworm.

### Hybrid genetic background of fall armyworm population invading China

In order to identify the specific genetic background of the invading fall armyworm samples in China, 105 samples from 16 provinces were re-sequenced, as well as four samples from two countries in Africa (Zambia and Malawi). A total of ~10 G genomic Illumina data were generated for each sample, with a total coverage of about 25 × genome size. Firstly, we analyzed the whole mitochondrial genome sequences of all samples. A total of 208 different SNP loci were screened based on the published mitochondrial sequences of both the American R-strain (AXE) and C-strain (ASW). The original Illumina reads of each sample were mapped to the reference mitochondrial genomes, and the genotype of re-sequenced samples were identified according to these SNP loci. We found that most of the samples were assigned to R-strain, and all four samples from Africa were C-strain, while only four out of 105 samples in China were assigned to C-strain based on the mitochondrial genome (Fig. 2a). The results were similar to those of the 173 Chinese fall armyworm samples identified by PCR based on the *COI* gene in this study. The proportion of C-strain was only about 10% (Supplementary Table 5), which is consistent with previous reports^42^.

**Figure 2.**
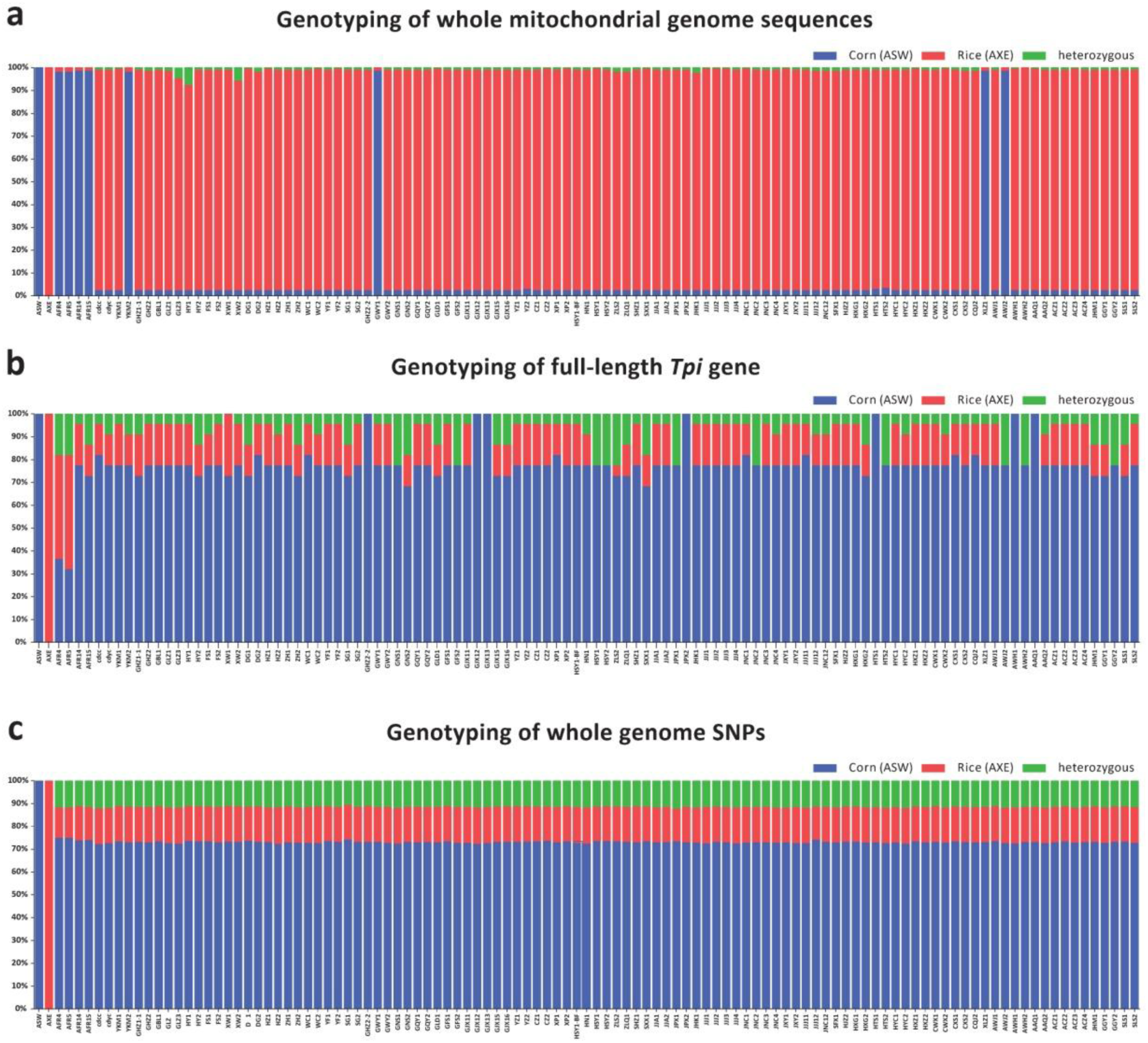
Genetic background of 109 fall armyworm samples based on two molecular markers and genome-wide SNPs. ASW represents the American corn strain and AXE represents the American rice strain. AFR4-5 were both from the inbred strain AFR2017 collected from Zambia, AFR14-15 were both from the same inbred strain AFR2019 collected from Malawi. cdcc and cdyc represent two inbred strains collected from Yunnan Province in China.

Next, we analyzed the *Tpi* gene, which is commonly used in strain identification^36–37^. By comparing the full length *Tpi* gene of published R-strain and C-strain fall armyworm, 22 SNP loci were found. The genotypes of all samples were analyzed by calculating the ratio of SNPs compared to the American reference sequences (AXE and ASW). The results showed that most fall armyworm samples collected from China contained more C-strain SNP loci, as did the Malawi samples (AFR14, AFR15), but not those from Zambia (AFR4, AFR5) which contained approximately 50% of R-strain SNP loci. Genotypes of seven samples were identical to the American C-strain (ASW) reference (Fig. 2b), possibly due to them being females or homozygous genotypes. The remainder of the samples contained a small proportion of R-strain genotypes or heterozygous SNPs. However, none of the samples were found to be identical to the American R-strain genotype (AXE). We designed a pair of primers to amplify a *Tpi* fragment that included *Tpi*-exon3, exon4 and intron3. We used the primers to screen 173 samples and to analyze 10 different SNP loci of each sample by comparing to R-strain and C-strain genotypes reported previously (Fig. 3, Supplementary Table 5). The results showed that almost all of the samples were C-strain genotypes except three samples (G-GXW11, G-GXW13, G-EP6) were identified as Africa-specific haplotype, in which 6 of 10 SNPs were identical to R-strain, while 4 were identical to C-strain, and 10 unique SNPs were significantly different from known R- or C-strain genotypes (Fig. 3). It can also be seen from the Figure 2b that the genotyping results of AFR4-5 based on full-length *Tpi* gene, which represents the Africa-specific strain, are quite different from the rest of the samples in containing >40% R-strain SNPs. In summary, our genotyping results show that there are obvious contradictions between strain identification using mitochondrial and nuclear *Tpi* gene markers.

**Figure 3.**
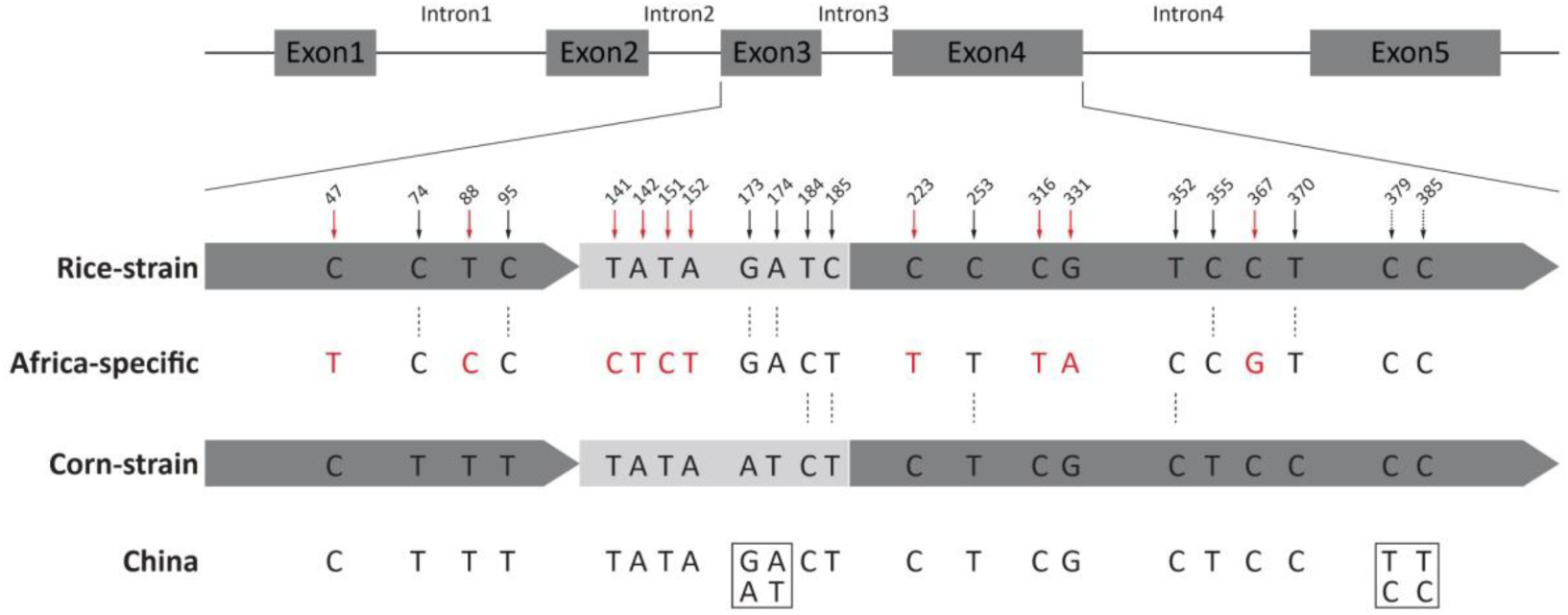
Diagram of the *Tpi* gene segments with respect to consensus Western Hemisphere sequences and the haplotypes observed in samples collected from Africa and China. Black solid arrows represent 10 SNPs used to identify traditional R-strain and C-strain fall armyworm, in which P370 was considered to be an effective diagnostic marker especially. Red solid arrows represent 10 SNPs specific to Africa-specific strain. The boxes represent two variable loci in some Chinese samples, including homozygous or heterozygous genotypes.

In order to clarify the genetic background of fall armyworm populations invading China, we mapped the published Illumina data of R-strain (AXE) and C-strain (ASW) to the reference genome of this study. Both AXE and ASW strains were from colonies that have been reared in laboratory for over 10 years, which could be considered to represent R-strain and C-strain populations, and finally a total of 707353 homozygous SNPs between the two strains were screened. The genotypes of 109 re-sequenced samples were analyzed by comparing Illumina data of each sample with these SNPs (Fig. 2c). The results showed that all the samples, including the four from Africa, had more than 70% of the genetic background of the American C-strain, but there were no individuals completely identical with C-strain (ASW) genotype. The proportion of R-strain SNPs was less than 15%, and the remaining 15% were heterozygous. The results showed that fall armyworm invading China have a dominant percentage of the C-strain background at the whole genome level. By comparing the results of the mitochondrial genome, *Tpi* gene and genome-wide identification, it becomes apparent that there is no obvious correlation between the mitochondrial and whole genome genotype. *Tpi* genotyping shows results more similar to those of the whole genome. However, there is a high proportion of heterozygosity at strain-specific loci, perhaps due to male individuals with ZZ chromosomes, which reduces the accuracy of genotyping based on the *Tpi* gene. Moreover, the presence of Africa-specific *Tpi-*haplotype increases the complexity of using this marker for identification. Strain analysis based on the whole genome reflects a more accurate approach.

In addition, 105 re-sequenced Chinese samples were collected from different regions of 50 cities distributed across 16 provinces. The collection time and sites coincided almost perfectly with the spreading invasion of fall armyworm in China. However, there was no obvious correlation between the time or site of collection and the genetic structure of the fall armyworm population (Fig. 2). Almost all samples have similar genomic backgrounds, which suggests that the invading population may originate from a single genetic source and there is no evidence for genomic selection during the invasion. However, the analysis based on the mitochondrial genome (Fig. 2a) and the *Tpi* gene (Fig. 2b) might erroneously suggest a more heterogeneous population genetic structure.

### Fall armyworm is developing high risk of resistance to conventional pesticides

Insecticide resistance evolution is one of the most challenging problems to be solved in the control of fall armyworm. Identifying resistance-related genes is helpful for the monitoring and prevention of fall armyworm outbreaks. We selected 14 previously reported resistance-related genes of lepidopteran pests (Supplementary Table 7) and scanned the re-sequenced samples to analyze variation in target genes. The results showed that all the target genes had multiple variation sites with high frequency of SNPs in the coding sequence (CDS) region (Fig. 4a). Sequence diversity of CYP450 and AChE were both reached 0.01, while the rest genes were no more than 0.005 (including heterozygotes), suggesting a great potential risk of resistance evolution.

**Figure 4.**
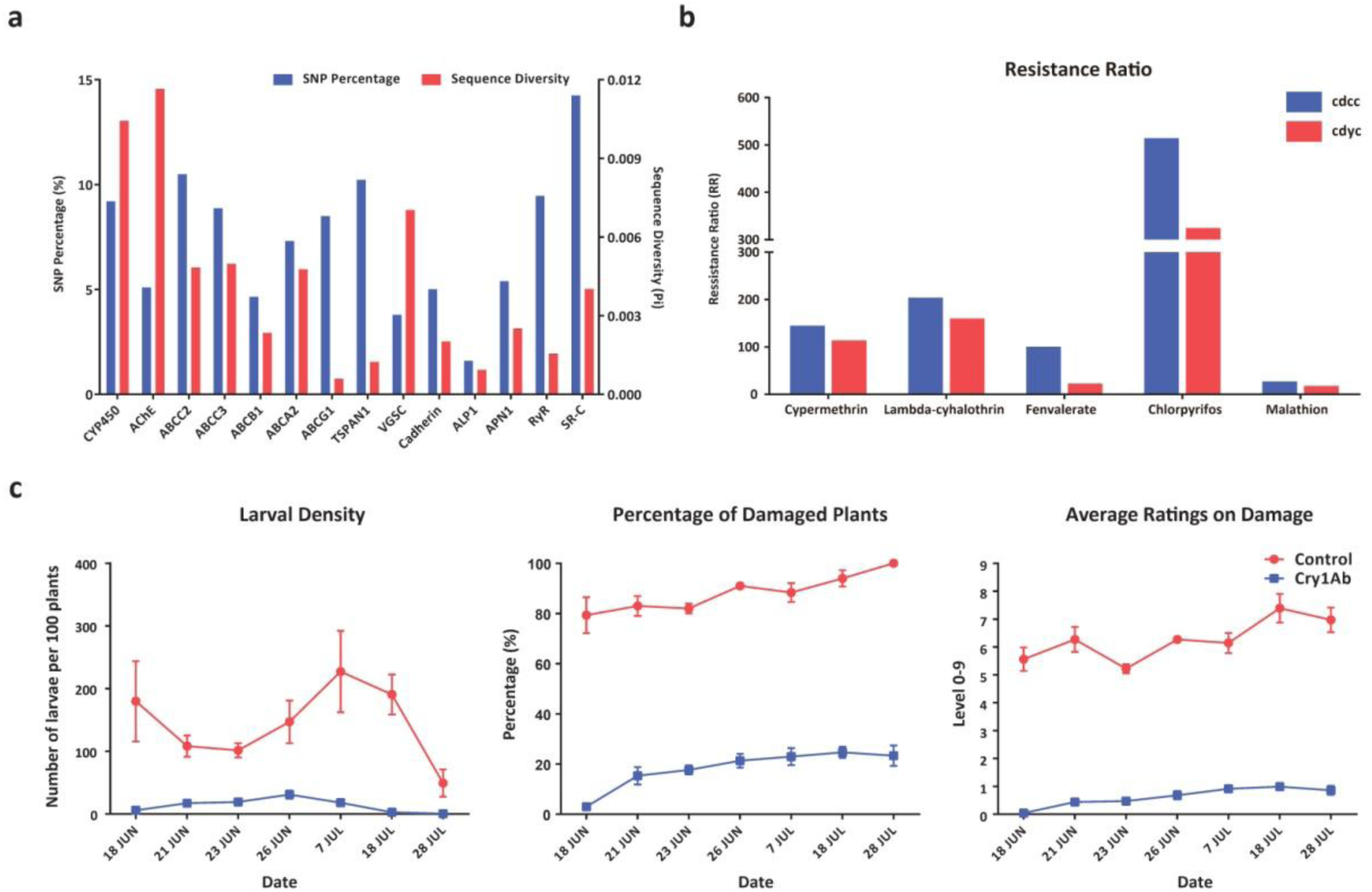
Genomic scanning and bioassays of fall armyworm for insecticides resistance. a) Sequence variation analysis of pesticides resistance and Bt resistance related genes in CDS region. SNP percentage means the proportion of variation sites in the complete gene CDS length. b) The resistance ratios (RRs) of two Chinese fall armyworm populations to pyrethroids (cypermethrin, lambda-cyhalothrin, fenvalerate) and organophosphates (chlorpyrifos, malathion) insecticides, The RRs was calculated by LD_50_ (µg/g) of field population over the LD _50_ of the susceptible population. The LD_50_s of the susceptible population here was referred to Yu et al. (1991). c) Resistance tests of GM maize and control maize to fall armyworm in field experiments.

Studies have shown that the amino acid substitutions in AChE (A201S, G227A, F290V), VGSC (T929I, L932F, L1014F) and RyR (I4790M, G4946E) result in resistance to organophosphate, pyrethroid and diamide insecticides, respectively. The results of variation scanning of the re-sequenced samples showed that resistance mutations were found at the first (AA201) and third locus (AA290) of AChE gene. Among them, the first locus had 17.1% heterozygous mutations, and the third locus had 29.7% homozygous resistance mutations and 58.2% heterozygous mutations. And no resistant mutations were detected at corresponding sites of VGSC and RyR gene in any samples (Table 1). We designed primers to detect the resistant mutation sites in *AChE* in 173 Chinese samples by PCR amplification and Sanger sequencing. The results were similar to the Illumina data, showing approximately 25% homozygous and 50% heterozygous variation at the third locus (AA290).

**Table 1.** Genotype and resistance mutation sites of pesticide-related genes in fall armyworm populations in China.

|  | Genotype |  |  |  |  |  |  |  |
| --- | --- | --- | --- | --- | --- | --- | --- | --- |
| Gene | AChE |  |  | VGSC |  |  | RyR |  |
| Mutation Sites | AA201 | AA227 | AA290 | AA929 | AA932 | AA1014 | AA4790 | AA4946 |
| Susceptible<br>(amino acid) | A | G | F | T | L | L | I | G |
| Resistance<br>(amino acid) | S | A | V | I | F | F | M | GE |
| Re-sequenced<br>Samples | AA<br>(82.9%)<br>AS<br>(17.1%) | GG<br>(100%) | FF<br>(12.1%)<br>FV<br>(58.2%)<br>VV<br>(29.7%) | TT<br>(100%) | LL<br>(100%) | LL<br>(100%) | II<br>(100%) | GG<br>(100%) |

We determined the LC_50_s to 14 insecticides for two Chinese fall armyworm populations collected from Yunnan Province (Supplementary Table 8). The results showed that the LC_50_s for both fall armyworm populations to fenvalerate, malathion, chlorpyrifos were relatively high. The LC_50_s of chlorantraniliprole and cyantraniliprole were low along with emamectin benzoate and E-MBI (Fig. 5). The resistance levels of the two populations to pyrethroids and organophosphate pesticides were very high; in particular, the resistance ratio to chlorpyrifos of both populations were more than 300-fold compared to a laboratory susceptible population that was sampled in 1975^52^ (Fig. 4b, Supplementary Table 9). These results provide a susceptible baseline for fall armyworm populations invading China to different pesticides, which can provide guidance for resistance monitoring and pesticide management strategies.

**Figure 5.**
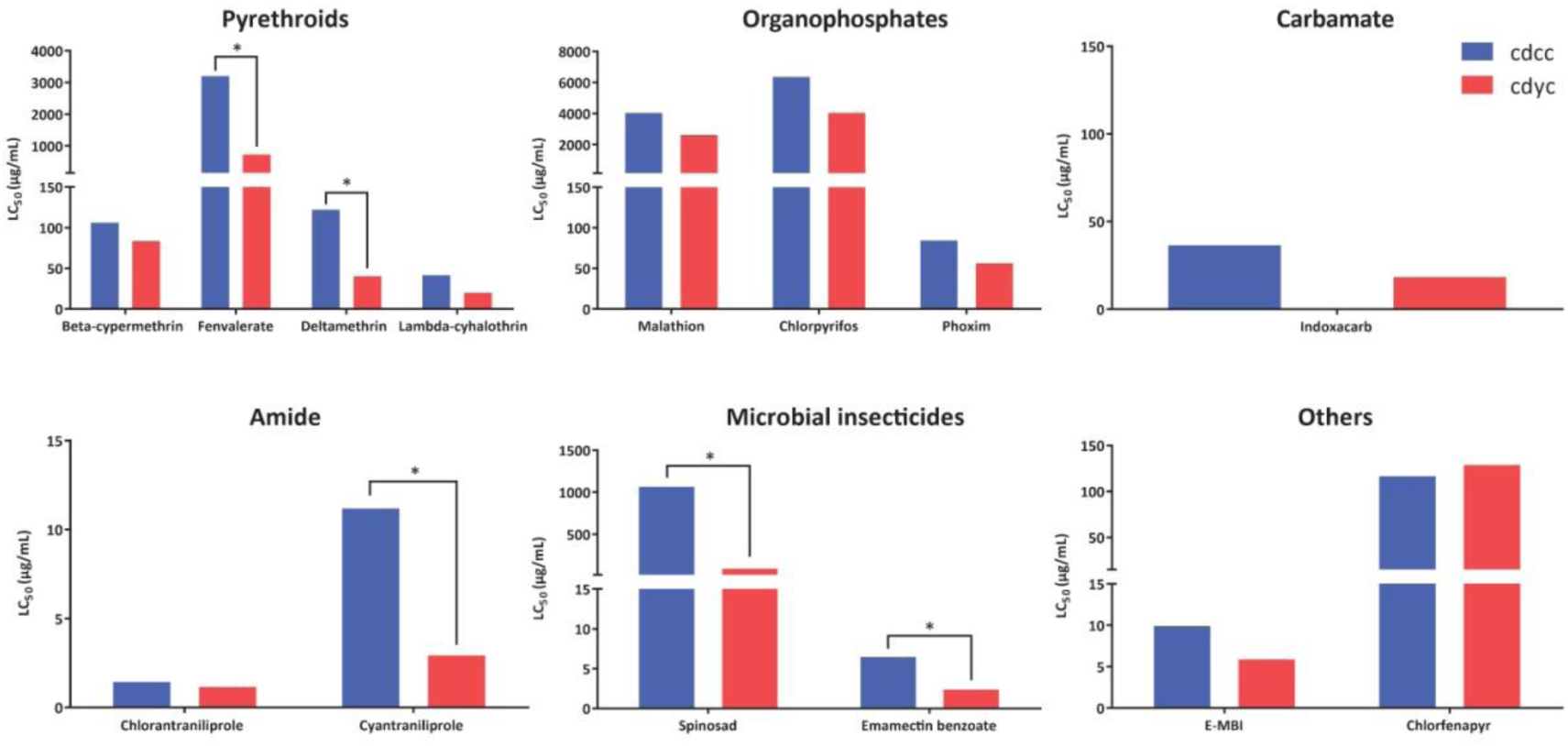
The LC_50_s of two Chinese fall armyworm populations to different kinds of insecticides. The significantly difference was considered by whether the 95% FL have overlap (*P<0.05).

### Fall armyworm invading China are currently sensitive to Bt toxin in field-evolved experiment

The insertion of 2 bp in the *ABCC2* of fall armyworm was reported to cause a frame-shift mutation and results in resistance to Cry1Fa^33^. We did not detect the same insertion mutation in 105 re-sequenced samples nor in 173 samples screened by using PCR and Sanger sequencing. Although the percentage of SNPs in the CDS region of other Bt receptors such as *SR-C* (scavenger receptor class C gene, a specific receptor for Vip3Aa in Sf9 cells), *TSPAN1* and other ABC gene-family related to Cry toxin were also very high (Fig. 4a), no reported resistant mutation were found in any target resistance genes.

The field tests showed that fall armyworm samples invading China were sensitive to GM maize expressing Cry1Ab compared with control group, which is accordant with the result of resistant genes scanning. The damage assessment on larval density, percentage of damaged plants and average damage ratings of GM maize were significantly lower than those of the control group (Fig. 4c), which indicated that the GM maize expressing Cry1Ab currently has good control effects on the invading population of fall armyworm in China.

## Discussion

The rapid spread of the fall armyworm has attracted popular attention worldwide. Accurate identification of its genetic characteristics (strain and pesticide resistance properties) has a direct and practical importance in terms of risk assessment and control strategies. A genome-wide analysis can reveal more in-depth genetic information than conventional gene-level analyses. The results of this study show that the fall armyworm invading China had a genetic background with dominant American corn-strain genotype and might be descendants of an inter-strain hybrid population. Most of the fall armyworm samples invading China were detected and collected from corn and sugarcane, which are more likely to show the characteristics of C-strain host plants. Along the invasion path of the migratory fall armyworm, there are large-scale rice planting areas in Southeast Asia and central China, however, so far there are few reports of serious damage to rice caused by fall armyworm (http://www.fao.org/fall-armyworm). The traditional R-strain fall armyworm in the Americas mainly feeds on turf grass, and there are also some reports of damage to rice. In addition, the traditional R-strain *Tpi* genotype has not been detected in any of the samples collected from Africa and Asia, so we speculate that the American R-strain fall armyworm did not invade Africa or Asia, including China.

Mitochondrial *COI* and *Tpi* genes are commonly used as markers for the identification of fall armyworm strain. Current studies have shown that the mitochondrial genome is clearly divided into two strains. Because of their hybridization, identification based on the maternally-inherited mitochondrial genotype is inaccurate and the insertion of two C-strain mitochondrial fragments in this study further confirms this inaccuracy. Therefore, it is not possible to accurately infer fall armyworm strain status based on the complete mitochondrial genome or the single *COI* mitochondrial gene, as suggested previously^40–41^. The *Tpi* gene, located in the nuclear genome, is more suitable for strain identification and 10 SNPs in this gene can distinguish between R- or C-strains fall armyworm in the Americas^37^. In this study, we found that the AT/GC SNP located at *Tpi*-intron3 (P173/174) was not linked with the other eight SNPs and was not specific to either the R- or C-strain genotypes precluding these two SNPs as diagnostic markers. In addition, the TT/CC SNP located at *Tpi*-exon4 (P379/385) was associated with sequence variation in *Tpi*-intron4 (Fig. 3, Supplementary Fig. 2). A special (Africa-specific) haplotype of the *Tpi* gene was identified in samples collected from southern-east Africa, which was also detected in some Chinese samples. For this specific haplotype, it is not identical to either American R- or C-strain genotypes, although it was tentatively designated as R-strain based on E4^183^ site (equal to P370 in Fig. 3 in this study) in previous studies^37^. Our genome-wide SNP analysis revealed that this haplotype contained more C-strain SNPs than R-strain.

This analysis shows that the fall armyworm shows a complicated population genetic structure. For example, the sample used for the genome sequencing in this study represents a combination of the special *Tpi* haplotype and C-strain *COI*. We also found combinations of the R-strain *COI* and special *Tpi* (sample G-XW13) as well as heterozygous forms of the *Tpi-*special and *Tpi-*C with the R-strain *COI* in two samples (G-GXW11, G-EP6). These combinations of different genotypes and extensive gene exchange show that the genetic boundaries between two traditional (American) R- and C-strains are obscure. The insertion of two C-strain mitochondrial fragments might be caused by random hybridization between different genotypes.

The rapid evolution of insecticide resistance and the increasing levels of resistance observed in fall armyworm populations needs attention. In this study, mutations related to organophosphate pesticide resistance were detected in AChE gene. Although some mutation sites were detected as heterozygous in most samples at present, the frequency of resistant mutation sites will increase greatly under the selection pressure caused by application of related pesticides in field. The pesticide bioassay results showed that armyworms invading China have evolved high levels of resistance to chlorpyrifos, in particular, which was consistent with the results of molecular scanning of insecticide resistance-related genes. However, the fall armyworms invading China are currently sensitive to GM maize expressing Cry1Ab in field experiments, and are also sensitive to other Bt toxins in the laboratory, according to previous studies^53^. At present, GM maize shows better application prospects in controlling fall armyworm in China, larval density and damage rate of GM maize were significantly less than that of normal plants, though this crop is currently not registered for use in the country.

This study provides a high-quality reference genome that demonstrates a genomic feature different from the traditional (American) C- or R-strain genotypes, as well as more comprehensive gene annotation. According to our results, commonly used strain identification of fall armyworms by mitochondrial or *Tpi* markers is limited or even inaccurate. We present resequencing data for 105 fall armyworm individuals invading China. The samples cover different regions and times during 2019, providing basic materials for global population genetic analyses. Baseline resistance data for Chinese fall armyworm populations is provided to 14 common pesticides, providing guidance for the control and resistance monitoring of fall armyworm. Small-scale field experiments in this study suggest that that fall armyworm are currently susceptible to GM maize, and these results could provide an important application reference for commercial planting of Bt maize in China. There are other important issues that remain for further exploitation using this whole genome approach, such as identifying the genes involved in polyphagy, migratory capability and olfaction, which could provide valuable tools for the future management of fall armyworms.

## Methods

### Genome sequencing and assembly

The fall armyworm samples were collected from maize fields in Lusaka, Zambia, in 2017 and reared to produce an inbred strain. One male moth, derived from seven successive generations of single-pair sib mating, was selected for extracting genomic DNA and constructing Illumina 350-bp and PacBio 20-kb insert libraries. Sequencing was performed on Illumina HiSeq 2500 and PacBio SMRT platforms, respectively. Two 3rd instar larvae were selected for Hi-C library construction and then sequenced on a HiSeq 2500 platform (Illumina). In addition, three 5th instar larvae, three pupae, three female moths and three male moths were used for RNA sequencing on an Illumina HiSeq 2500 sequencer. All the above samples were from the same inbred strain.

The raw PacBio reads longer than 5 Kb were assembled into contigs using software wtdbg2^45^. Arow v2.1.0 software was used to correct assembly errors after comparing contigs with PacBio reads by pbalign v0.4.1. The Illumina raw reads were filtered by trimming the adapter and low-quality regions, resulting in high-quality reads. Then clean Illumina reads were aligned to the assembled contigs by BWA v0.7.17, and single base errors in the contigs were corrected by Pilon v1.21.

After removing adapter and low-quality sequences, Hi-C sequencing data were aligned to assembled contigs with Hic-Pro, then the unmapped paired, dangling paired, self-circle and dumped reads were filtered out (https://github.com/nservant/HiC-Pro). According to the restriction enzyme (DpnII) sites on contigs, we clustered the linkage group of contigs based on agglomerative hierarchical clustering method by using LACHESIS^54^ (https://github.com/shendurelab/LACHESIS), and then these contigs were clustered into chromosomes.

### Gene annotation and prediction

A *de novo* repeat library of fall armyworm was constructed by RepeatModeler v. 1.0.4. TEs were identified by RepeatMasker v4.0.6 using both *de novo* library and Repbase library, and Tandem repeats were predicted using Tandem Repeats Finder^55^ v4.07b. The gene models were predicted by EVidence Modeler^56^ v1.1.1, combined with *ab initio* predictions, homology-based searches, and RNA sequencing alignments. Predicted gene models supported by at least one of the annotations using UniProt datbase, NR database, and RNA-seq data were retained. Then gene functional annotation was performed by aligning the protein sequences to the NCBI NR, UniProt, eggNOG, and KEGG databases with BLASTp v2.3.0+.

### Phylogenetic tree construction and genomic comparison

Orthologous and paralogous groups of 10 species (*Drosophila melanogaster*, *Plutella xylostella*, *Bombyx mori*, *Manduca sexta*, *Danaus plexippus*, *Heliconius melpomene*, *Operophtera brumata*, *Helicoverpa armigera*, *Spodoptera frugiperda*, *Spodoptera litura*) with published genomes were analyzed by OrthoFinder v2.3.1 with default parameters. Orthologous groups that contain single-copy genes for each species were selected to construct the phylogenetic tree. The multi-sequence alignment of proteins was accomplished by MUSCLE^57^ v3.8.31. The Neighbor-Joining (NJ) phylogenetic tree was constructed by MEGA v7.0.14. The current assembled genome was aligned with two published versions of fall armyworm genomes by nucmer software with cutoff of identity >80% and coverage >80%. Unique alignment comparison was conducted with the parameters "delta-filter -i 85 -l 10 -r -q", while multiple alignment comparison were conducted with the parameters "delta-filter -i 85 -l 10".

### Sampling for resequencing and population genetic study

A total of 105 Chinese fall armyworm samples were used for resequencing, including four samples of two inbred strains (cdcc and cdyc) collected from Yunnan Province and reared for multiple generations in the laboratory. All samples were collected as larvae on maize or sugarcane from 50 cities of 16 provinces (autonomous regions or municipalities) of China. The larvae were fed with fresh maize leaves and brought back to the laboratory under ambient conditions during transportation. Larval bodies were cleaned and then stored in a freezer at −80 ^o^C. The detailed sample information is shown in Supplementary Table 10 and the sample distribution in China is shown in Supplementary Figure 3. In addition, four fall armyworm samples from Africa were also used for resequencing, including two samples (AFR4-5) from the same inbred strain (AFR2017) as the genome sequencing in this study, and another two samples (AFR14-15) which were collected from maize fields in Bvumbwe, Malawi, in January 2019, which is also an inbred strain (AFR2019) reared in laboratory. After DNA extraction of each sample, a 350-bp insert library were constructed and paired-end sequencing was performed following the standard Illumina protocol. Sequence reads from microorganisms and host plants in raw data were removed before analysis.

In addition to re-sequenced samples, another 173 fall armyworm samples from 21 provinces in China were used for strain identification and molecular detection based on PCR amplification and Sanger sequencing. The samples were collected from the field as larvae or adult moths. The detailed sample information is shown in Supplementary Table 5 and the sample distribution in China is shown in Supplementary Figure 3. Mitochondrial *COI* and *Tpi* markers were used for strain identification. ABCC2, AChE genes were detected based on primers designed according to published mutation sites^31–33^. Mitochondrial insertion fragment detection was conducted using primers designed in this study. All primer sequence information in this study is shown in Supplementary Table 11.

### Read mapping and SNP calling

The Illumina raw reads from re-sequenced samples were filtered using clean_adapter and clean_lowqual software (https://github.com/fanagislab/common_use), resulting in high-quality reads with an average error rate of < 0.01. Then, the high quality reads were aligned to the fall armyworm reference genome and mitochondrial genome sequences by the Burrows-Wheeler Transform Alignment (BWA) software^58^ package v0.7.5a with default parameters. Alignments for each sample were processed by removing duplicate reads using samtools^59^ software package v1.3. The mpileup function in samtools was used to generate mpileup files for each sample. Bcftools-vcftools^60^ was used to identify SNPs and small Indels. Several criteria were considered in SNP filtering: (1) a read mapping score higher than 40; (2) minimum coverage greater than 10.

### Bioassays of insecticides and Bt maize in the field

Bioassays were conducted by a topical application procedure^61^. Two inbred strains of Chinese fall armyworm populations (cdcc, cdyc) were tested using 14 types of pesticide commonly used in agricultural production (Supplementary Table 12). 1.0 µL drops of serial dilution of technical insecticides in acetone solution were applied with a micropipette to the thoracic dorsum of the 3rd instars and the control larvae were treated with 1.0 µL acetone. After treatment, the larvae were reared individually in 24-well plates containing ad libitum artificial diet without any Bt proteins and insecticides. Larvae were retained in an insect chamber with a controlled environment of 26 ± 1°C, 60 ± 10% RH and a photoperiod of 16 h: 8 h (L: D). Mortality was assessed after 72 h treatment. Larvae were considered dead if they were unable to move in a coordinated manner when prodded with a small soft brush. We used median lethal doses LC_50_ to evaluate the resistance level of different fall armyworm populations. The LC_50_ for each assay of insecticides was estimated by probit analysis using the software package POLO-PC^62^ (LeOra Software, Berkeley, CA, USA).

The Bt toxin field bioassays to were conducted at a genetically modified (GM) test base in Yunnan Province, China. Test seeds of GM maize (expressing Cry1Ab) and control maize were provided by DBN Biotech Center, Beijing DBN Technology Group Co., Ltd. Both maize types were planted in approximately 180 m^2^, with each type being replicated three times. Larval density and maize damage rates were investigated at different growth stages of maize at seven different dates during June to July. The investigation was performed in a five-spot-sampling method with 20 maize plants per point. Fall armyworm damage assessment was performed according to standard procedures^63–65^.

## Acknowledgements

The following bodies provided funding that contributed to this work: Key Project for Breeding Genetic Modified Organisms grant (2016ZX08012004-003), the UK’s Global Challenges Research Fund and Biotechnology and Biological Sciences Research Council (BB/P023444/1), the UK Natural Environment Research Council Envision Doctoral Training Programme (NE/L002604/1).

---

**Supplementary Figure 1.**
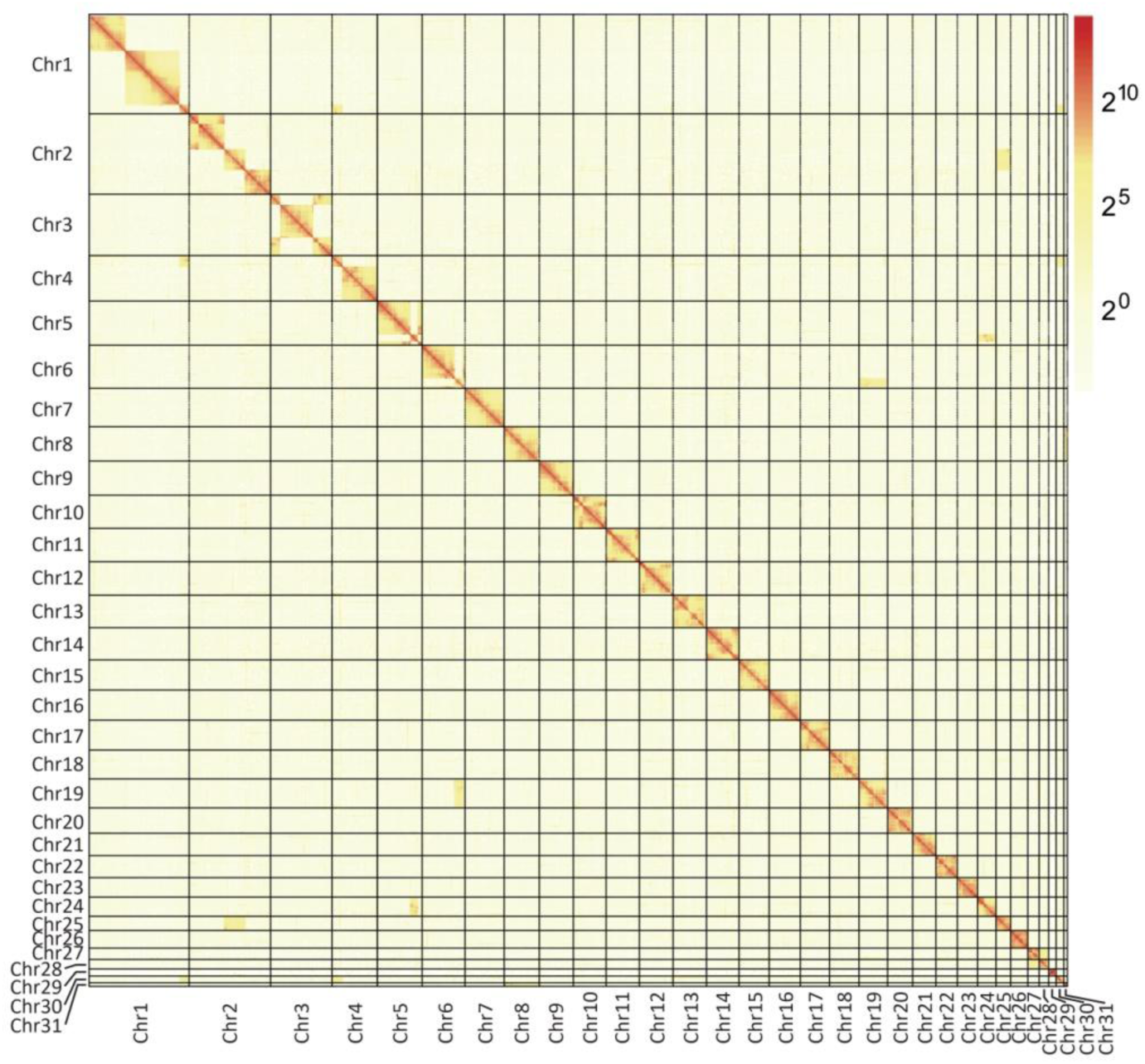
A genome-wide contact matrix from Hi-C data between each pair of the 31 chromosomes. The color value indicates the number of valid reads.

**Supplementary Figure 2.**
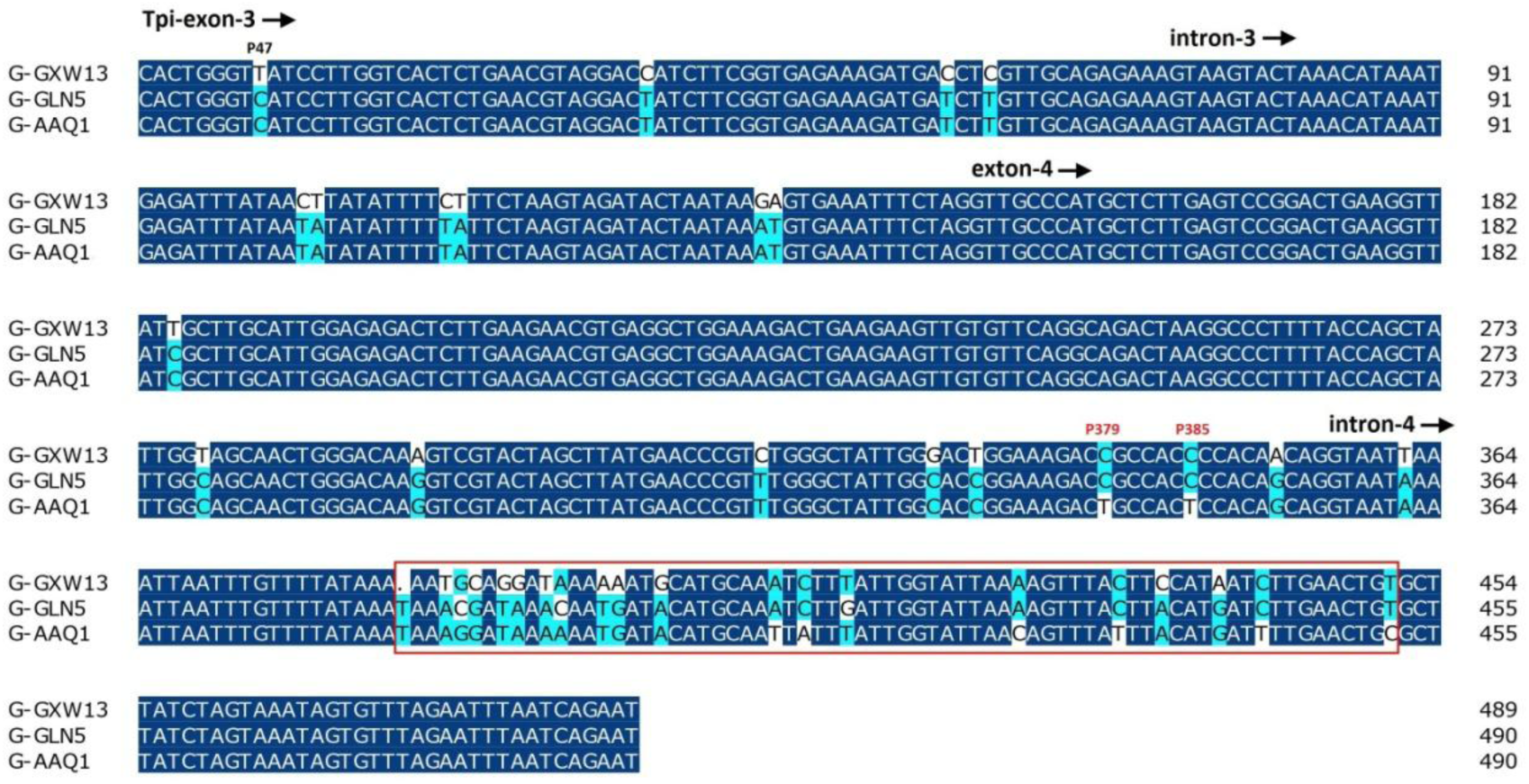
Comparison of different Tpi genotypes in Chinese samples. G-GXW13 represents Africa-specific genotype, G-GLN5 and G-AAQ1 represent C-strain genotype with difference in intron4 region. Red box represents the sequence variation in intron4 region linked to two variable sites (P379/385) in exon-4 of Tpi gene.

**Supplementary Figure 3.**
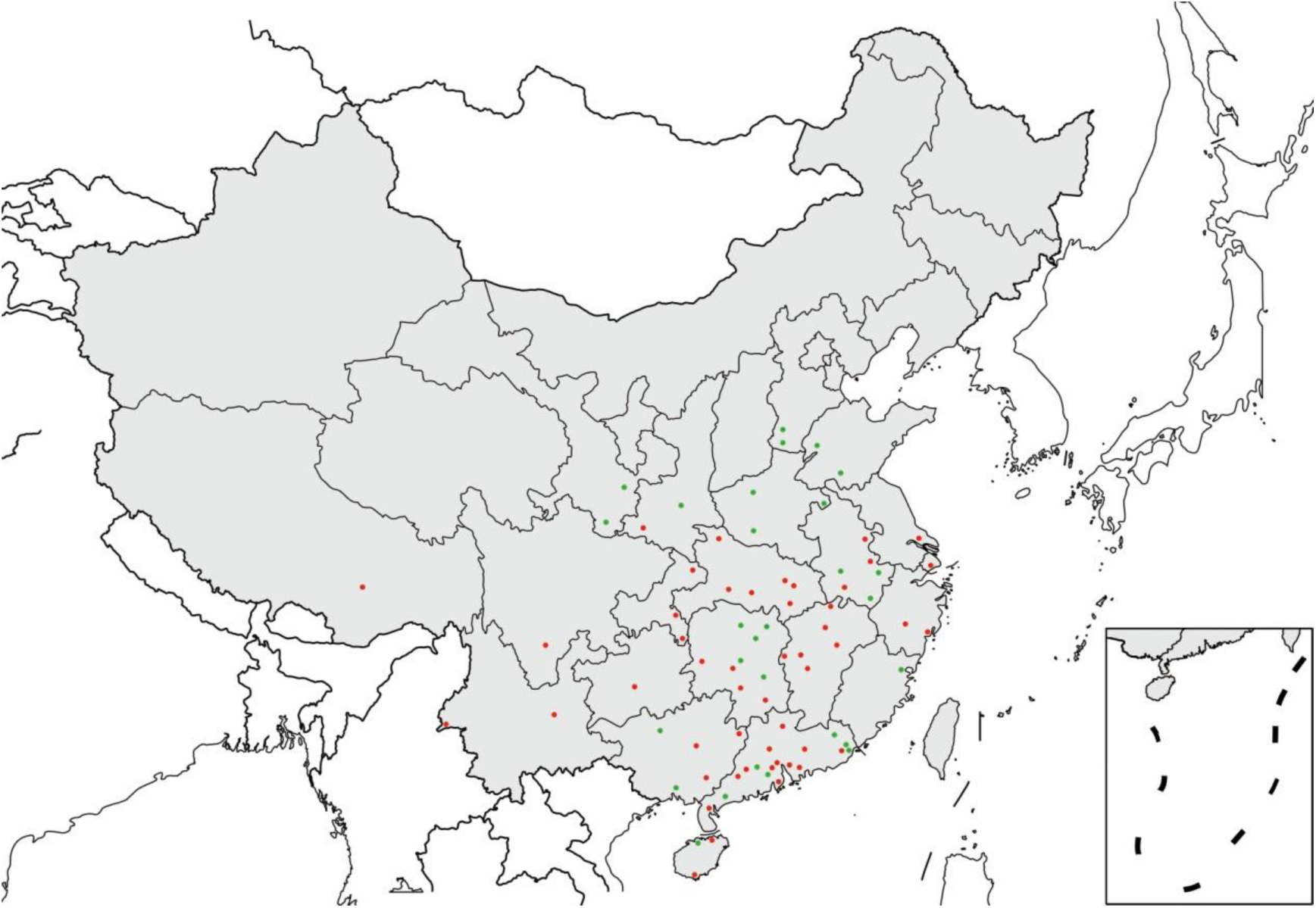
Map of fall armyworm collection sites in China. Each point represents a city, the red point represents the source of the re-sequenced samples from 16 Provinces in China, the green points represents the source of some rest samples from other 5 Provinces used for population genetic study in this study.

**Supplementary Table 1. Statistical results of chromosome lengths by Hi-C assembly.**

**Supplementary Table 2. Comparison between two versions of published fall armyworm genomes and the genome of this study.**

**Supplementary Table 3. Annotation and distribution of repetitive elements in fall armyworm genome.**

**Supplementary Table 4. Summary of assembly and annotation of Lepidoptera and Diptera genomes.**

**Supplementary Table 5. Information of 173 fall armyworm samples used for population genetic analysis in this study.**

**Supplementary Table 6. Distribution of mitochondrial insertion in 109 re-sequenced fall armyworm samples and American corn-strain (ASW) and rice-strain (AXE).**

**Supplementary Table 7. Genes related to pesticide resistance and Bt resistance in fall armyworm.**

**Supplementary Table 8. The LC_50_s of two Chinese fall armyworm populations to 14 kinds of insecticides.**

**Supplementary Table 9. The resistance ratios of two Chinese fall armyworm populations to five kinds of pyrethroids and organophosphorus insecticides.** The resistance ratio was calculated by LD_50_ (µg/g) of field population over the LD _50_ of the susceptible population.

**Supplementary Table 10. Information of 109 re-sequenced fall armyworm samples in this study.**

**Supplementary Table 11. Primer sequence information used in this study.**

**Supplementary Table 12. General information of 14 kinds of insecticides used in topical bioassays.**

## Notes

#### Summary of Updates

Figure 4b revised; Table 1 revised.

